# Targeting PI3Kβ-dependent cancer with a novel small molecule inhibitor, GT220

**DOI:** 10.64898/2026.01.16.699992

**Authors:** Qi Wang, Huimin Cheng, Xiangyang Yu, Changli Qian, Renlei Ji, Thomas M. Roberts, William D. Kerns, Jean J. Zhao

## Abstract

PI3Kβ is a critical oncogenic driver in cancers harboring PTEN loss or PIK3CB alterations, yet effective and selective PI3Kβ-targeted therapies remain elusive. Here, we report the development and preclinical characterization of GT220, a highly selective and potent small-molecule PI3Kβ inhibitor developed through integrated artificial-intelligence–driven design with medicinal chemistry and pharmacologic optimization. GT220 exhibits exceptional biochemical selectivity for PI3Kβ, binding with sub-nanomolar affinity and with minimal activity against other class I PI3K isoforms or the broader protein kinome. In cellular models, GT220 potently suppresses AKT phosphorylation and selectively inhibits viability of PTEN-deficient cancer cells, while sparing PTEN/PIK3CB wild-type and PI3Kα-dependent cells. In vivo, GT220 achieves a favorable tumor exposure with sustained PI3Kβ pathway inhibition and demonstrates robust antitumor efficacy and good tolerability in PTEN-deficient breast cancer xenograft models. In contrast, GT220 shows no antitumor activity or pathway inhibition in PTEN-wild-type or PI3Kα-dependent tumors, underscoring its context-dependent mechanism of action. Collectively, these findings establish GT220 as a promising next-generation PI3Kβ inhibitor and provide a strong preclinical rationale for precision targeting of PI3Kβ-dependent cancers.

## Introduction

Phosphatidylinositol-3,4,5-triphosphate (PIP3) is a membrane lipid molecule that plays a crucial role in cellular signaling by activating the phosphatidylinositol 3-kinase (PI3K) pathway. This regulates cell proliferation, motility, and survival. The levels of PIP3 are tightly regulated by the opposing actions of PI3K which phosphorylates phosphatidylinositol-4,5-bisphosphate (PIP2) to generate PIP3, and a lipid phosphatase PTEN (phosphatase and tensin homolog), which dephosphorylates PIP3 back to PIP2. Dysregulation of these enzymes can lead to excessive accumulation of PIP3, resulting in hyperactivation of the PI3K pathway and promoting cell proliferation and cancer development [1, 2].

Class I PI3Ks are subdivided into class IA (PI3Kα, β, and δ), which function as heterodimers composed of a p110 catalytic subunit and a p85 regulatory subunit and are activated downstream of receptor tyrosine kinases (RTKs) and other signaling inputs, and class IB (PI3Kγ), which is primarily regulated by G-protein–coupled receptors. Among these class 1A isoforms, PI3Kα and PI3Kβ are ubiquitously expressed, whereas PI3Kδ and PI3Kγ are largely restricted to hematopoietic cells. The distinct regulatory mechanisms, tissue expression patterns, and genetic dependencies of PI3K isoforms have created opportunities for isoform-selective therapeutic targeting aimed at improving efficacy while minimizing systemic toxicity [1, 2].

Oncogenic alterations in *PIK3CA*, encoding PI3Kα, represent one of the most common driver events in cancer, particularly in breast and endometrial tumors [3, 4]. This understanding has led to the successful clinical development of PI3Kα-selective inhibitors, including alpelisib and, more recently, inavolisib, both of which have demonstrated clinical benefit in biomarker-defined patient populations [5, 6]. By comparison, although somatic *PIK3CB* mutations are less frequent, they are recurrent across multiple cancer types and are increasingly recognized as functionally significant oncogenic events [7-9]. Despite this, no approved therapies currently target *PIK3CB*-altered tumors, and the development of selective PI3Kβ inhibitors has lagged behind that of other PI3K isoforms.

Loss of PTEN function represents another major mechanism of PI3K pathway hyperactivation and is among the most common alterations across human cancers [10, 11]. Beyond genetic loss-of-function mutations, PTEN activity is frequently suppressed through epigenetic, transcriptional, or post-translational mechanisms, resulting in functional PTEN deficiency in up to 30–80% of advanced tumors, depending on cancer type. Although PTEN-deficient cancers were initially expected to respond broadly to PI3K inhibition, early clinical trials with pan-PI3K and PI3Kα-selective inhibitors yielded disappointing results [12-14].

Through a series of foundational studies, our laboratory and others have demonstrated that these outcomes reflect a critical isoform dependency: tumors driven by RTK signaling or oncogenic *PIK3CA* mutations rely predominantly on PI3Kα, whereas PTEN-deficient tumors are uniquely dependent on PI3Kβ for survival and growth [15-19]. Since our initial discovery of this dependency [15], PI3Kβ has been validated as a key driver of oncogenesis in PTEN-null cancers across multiple preclinical models [20-23]. More recently, PI3Kβ has also been implicated in immune evasion programs in PTEN-deficient tumors [24] and in resistance to PI3Kα-targeted therapies, further underscoring its therapeutic relevance [23, 25]. Importantly, PI3Kβ appears to be largely dispensable in most adult tissues, supporting the possibility of an improved therapeutic index for PI3Kβ-selective inhibition.

Despite this strong biological rationale, initial PI3Kβ inhibitors encountered significant limitations. Early compounds such as SAR260301 were discontinued due to inadequate pharmacokinetics and efficacy [26], whereas others, including GSK2636771 and AZD8186, suffered from dose-limiting toxicities likely arising from insufficient potency or off-target inhibition of additional PI3K isoforms, particularly PI3Kδ [27, 28]. These challenges highlight the unmet need for PI3Kβ inhibitors with improved potency, selectivity, tumor exposure, and tolerability.

Here, we report the discovery and preclinical characterization of GT220, a highly selective and potent small-molecule PI3Kβ inhibitor developed through integrated artificial-intelligence–driven design with medicinal chemistry and pharmacologic optimization. We demonstrate that GT220 exhibits exceptional biochemical selectivity for PI3Kβ, potently suppresses AKT signaling and cell viability in PTEN-deficient and *PIK3CB*-mutant cancer cells, and achieves a robust tumor exposure with sustained pathway inhibition in vivo. Importantly, GT220 shows strong antitumor activity in PTEN-deficient xenograft models while sparing PTEN-wild-type and PI3Kα-dependent tumors. Together, these findings position GT220 as a promising next-generation PI3Kβ inhibitor and provide compelling support for precision targeting of PI3Kβ-dependent cancers driven by PTEN loss or oncogenic *PIK3CB* alterations.

## Results

### GT220 is a highly selective PI3Kβ inhibitor with potent biochemical activity

GT220 was developed through an integrated small-molecule discovery and optimization campaign in which approximately 1.2 million compounds were evaluated using artificial-intelligence–driven design (AIDD), computer-aided drug design (CADD), and medicinal chemistry approaches. This iterative process incorporated compound synthesis, in vitro PI3Kβ activity assays, and pharmacokinetic (PK) and pharmacodynamic (PD) profiling, ultimately leading to the selection of GT220 as a lead PI3Kβ inhibitor (Figure 1A).

**Figure 1.**
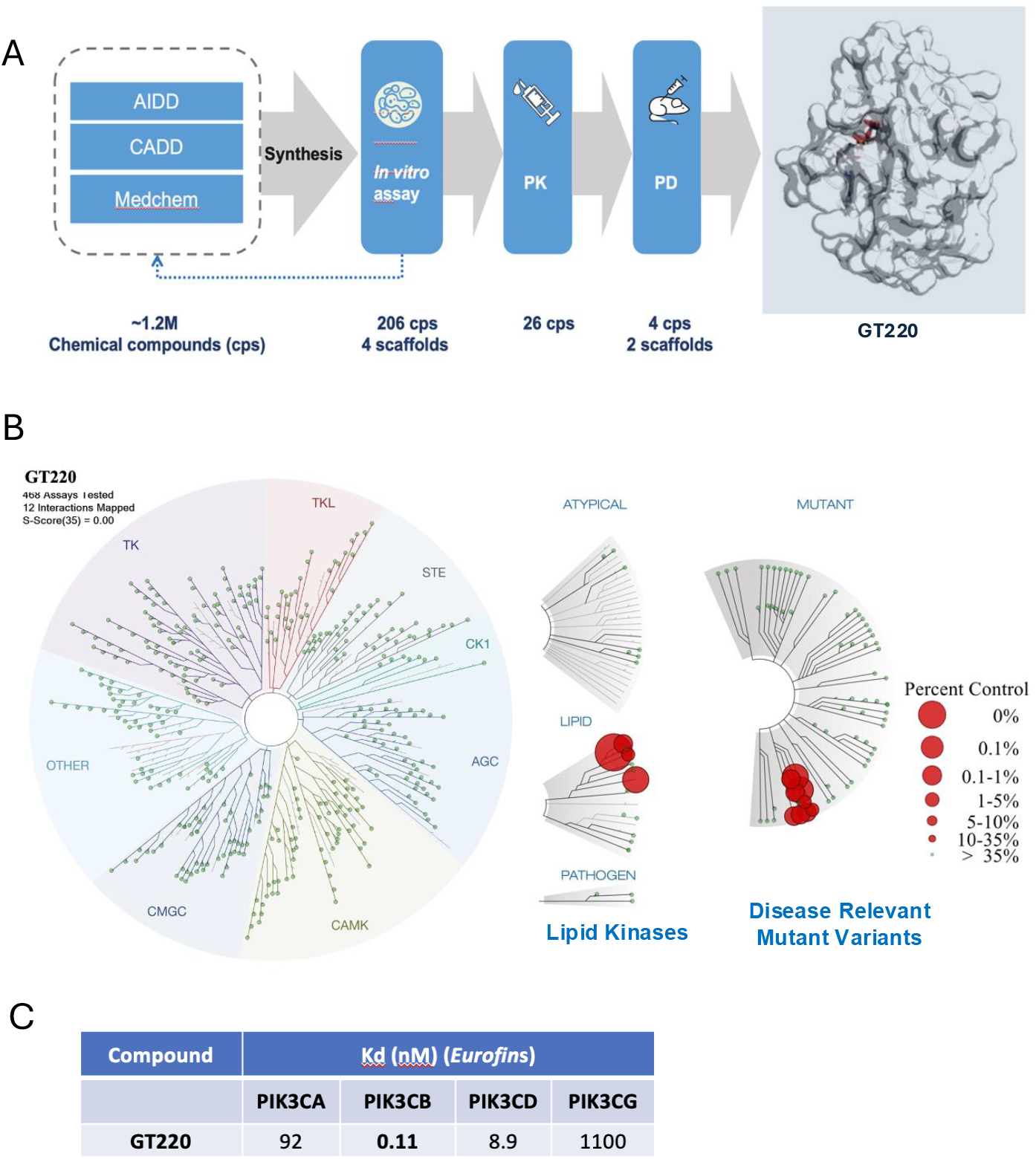
Development of the p110β-selective inhibitor GT220. **(A)** Schematic overview of the GT220 discovery and optimization workflow. Approximately 1.2 million compounds were evaluated through an integrated artificial-intelligence–driven design (AIDD), computer-aided drug design (CADD), and medicinal chemistry pipeline, followed by iterative synthesis, in vitro PI3Kβ activity assays, pharmacokinetic (PK), and pharmacodynamic (PD) profiling, leading to the identification of GT220 as a lead compound. **(B)** Kinome-wide selectivity profiling of GT220 assessed using the KinomeScan platform (Eurofins). GT220 was screened against a broad panel of human kinases, demonstrating high selectivity for lipid kinases and minimal off-target engagement across the kinome. Expanded views highlight selectivity within lipid kinases and disease-relevant mutant kinase variants, with percent control values indicated by node size and color. **(C)** Biochemical binding affinities of GT220 for class I PI3K isoforms measured using recombinant human proteins (Eurofins). Dissociation constants (K_d, nM) demonstrate strong selectivity for PI3Kβ (PIK3CB) relative to PI3Kα (PIK3CA), PI3Kδ (PIK3CD), and PI3Kγ (PIK3CG).

The kinase selectivity profile of GT220 was subsequently assessed using the KinomeScan platform, which measures active site–directed binding across 468 human kinases, including disease-relevant mutant variants, at a screening concentration of 10 µM. Strikingly, GT220 exhibited exceptional selectivity for lipid kinases, with no detectable binding to the protein kinase family (Figure 1B). Within the lipid kinase subset, *PIK3CB* (PI3Kβ) emerged as the top-ranked target. Limited binding was observed to a small number of mutant PI3Kα variants, including PIK3CA-E545K and PIK3CA-Q546K, at the screening concentration, without evidence of broad PI3Kα engagement (Figure 1B). These findings indicate that GT220 is not only PI3Kβ-selective, but also highly specific across the broader kinome.

To quantitatively define isoform selectivity, binding affinities of GT220 for class I PI3K isoforms were measured using the KinomeScan KdELECT assay. GT220 bound PI3Kβ with sub-nanomolar affinity (K_d_ ≈ 0.11–0.15 nM), while exhibiting at least ∼80-fold weaker binding to PI3Kα, PI3Kδ, and PI3Kγ (Figure 1C). Together, these biochemical data establish GT220 as a highly potent and selective PI3Kβ inhibitor with exceptional kinome specificity.

### GT220 selectively inhibits AKT signaling and cell viability in PTEN-deficient cancer cells

PI3Kβ signaling has been shown to drive AKT activation in tumors harboring *PTEN* loss [21, 25]. To evaluate the cellular activity of GT220, we first examined its ability to suppress AKT phosphorylation in PTEN-deficient cancer cells and compared its activity with established PI3Kβ inhibitors, including KIN193 (also known as AZD6482) [25], AZD8186 (a dual PI3Kβ/δ inhibitor) [28, 29], and GSK2636771 [27], which have been reported to target PI3Kβ-dependent tumors. In PTEN-null HCC70 cells, which lack *PTEN* and rely on PI3Kβ for signaling, a 1-hour treatment with GT220 resulted in a marked reduction of AKT phosphorylation at both Thr308 and Ser473 (Figure 2A). Quantitative analysis showed that although all comparator PI3K inhibitors significantly reduced phospho-AKT levels relative to control, GT220 achieved a greater degree of AKT inhibition at the tested concentration (Figure 2A).

**Figure 2.**
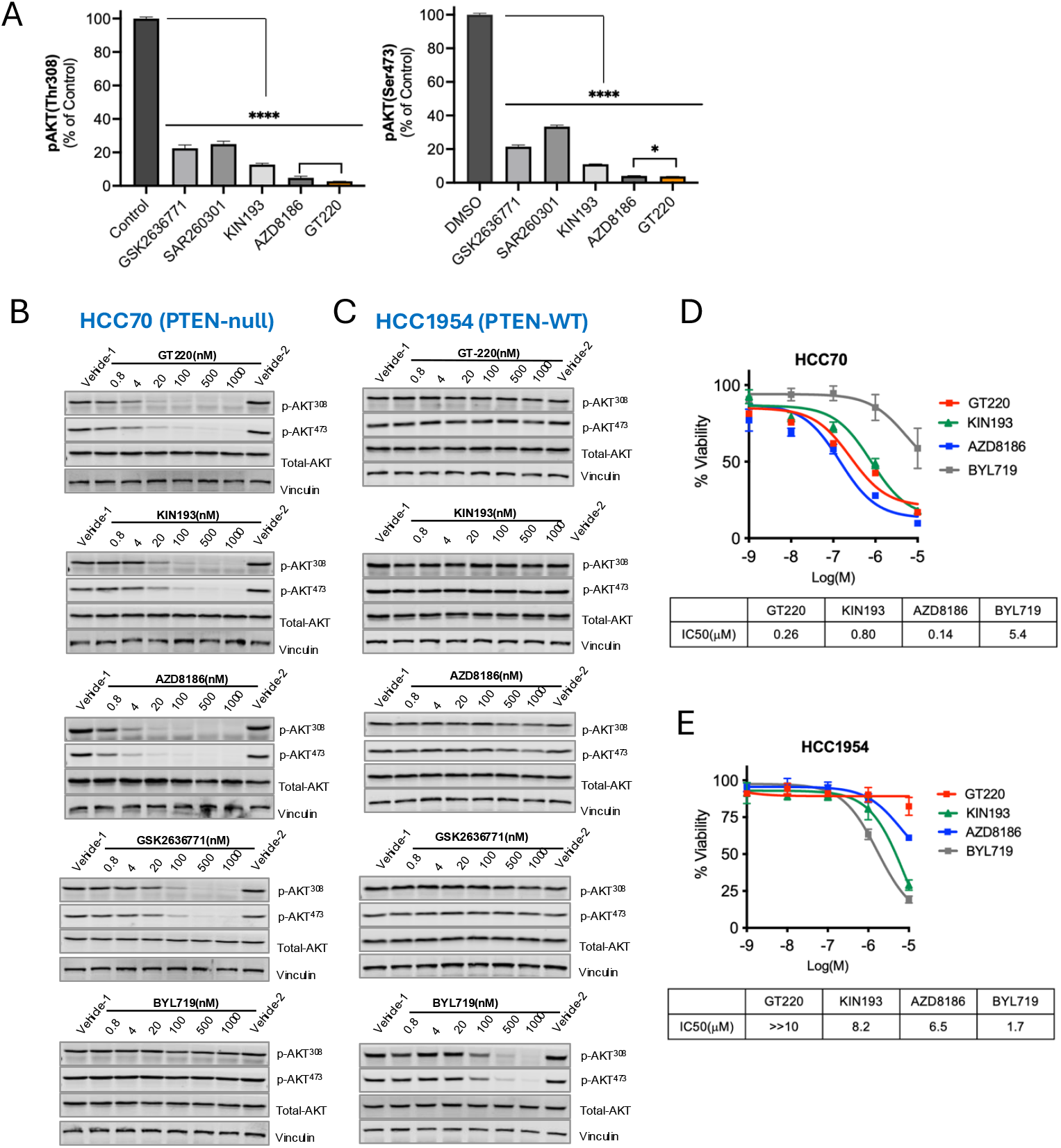
Selective inhibition of AKT signaling and growth by GT220 in PTEN-deficient cancer cells. **(A)** Quantification of AKT phosphorylation at Thr308 and Ser473 in PTEN-null HCC70 cells following 1-hour treatment with GT220 or the indicated PI3K inhibitors (0.5 μM). Phospho-AKT levels are expressed as a percentage of control (mean ± SD). Statistical significance is indicated (P < 0.05; ****P < 0.0001). **(B-C)** Immunoblot analysis of AKT signaling in HCC70 (PTEN-null; **B**) and HCC1954 (PTEN wild-type; **C**) cells treated for 1 hour with increasing concentrations of GT220, KIN193 (PI3Kβ-selective), AZD8186 (PI3Kβ/δ), GSK2636771 (PI3Kβ-selective), or BYL719 (PI3Kα-selective). Phosphorylation of AKT at Thr308 and Ser473, total AKT, and vinculin (loading control) are shown. **(D-E)** Cell viability dose–response curves for HCC70 (**D**) and HCC1954 (**E**) cells treated for 72 hours with increasing concentrations of GT220 or comparator PI3K inhibitors. Cell viability is expressed relative to vehicle-treated controls (mean ± SD). Tables summarize IC_50_values (µM) derived from nonlinear regression analysis.

The effects of GT220 were compared with established PI3K inhibitors in a pair of cancer cell lines representing distinct PI3K dependencies: PTEN-deficient, PI3Kβ-dependent HCC70 cells and PTEN-proficient, *PIK3CA*-mutant, PI3Kα-dependent HCC1954 cells. In PTEN-null HCC70 cells, GT220 induced strong, dose-dependent suppression of AKT phosphorylation at both Thr308 and Ser473, comparable to or exceeding that observed with the PI3Kβ-selective inhibitors KIN193 and GSK2636771, as well as the dual PI3Kβ/δ inhibitor AZD8186 (Figure 2B). In contrast, the PI3Kα-selective inhibitor BYL719 had minimal effect on AKT phosphorylation in HCC70 cells, even at higher concentrations, consistent with PI3Kβ-dependent signaling in this model.

In PTEN–WT HCC1954 cells, which harbor an activating *PIK3CA* mutation and rely on PI3Kα signaling, GT220 and other PI3Kβ inhibitors failed to suppress AKT phosphorylation across the tested dose range (Figure 2C). As expected, BYL719 effectively reduced AKT phosphorylation in these cells, confirming pathway specificity and appropriate cellular context dependence.

To determine whether pathway inhibition translated into functional growth effects, we assessed cell viability following 72 hours of treatment. GT220 potently reduced the viability of PTEN-null HCC70 cells in a dose-dependent manner, with IC_50_ values comparable to or lower than those of other PI3Kβ-targeting agents (Figure 2D). In contrast, GT220 had minimal effects on the viability of PTEN– wild-type HCC1954 cells, even at the highest concentrations tested (10μM), unlike KIN193 and AZD8186, which reduced viability in this context (Figure 2E). These findings indicate that GT220 exhibits high selectivity for PI3Kβ-dependent cells and spares PTEN–wild-type cells even at elevated concentrations, consistent with a targeted response.

Collectively, these results demonstrate that GT220 selectively suppresses AKT signaling and cell growth in PTEN-deficient cancer cells, while sparing PI3Kα-dependent, PTEN–wild-type cells— consistent with its biochemical selectivity for PI3Kβ.

### GT220 achieves favorable tumor exposure and robust PI3Kβ pathway inhibition in PTEN-deficient xenografts

To further characterize the in vivo pharmacodynamic (PD) and pharmacokinetic (PK) properties of GT220 relative to other PI3Kβ inhibitors, we conducted short-term dosing studies in PTEN-deficient HCC70 xenograft–bearing mice. Tumor-bearing BALB/c nude mice were treated orally twice daily with GT220, KIN193, or AZD8186 at doses of 30, 60, or 90 mg/kg for 3 days. Tumors and plasma were harvested 1.5 hours after the final dose for biochemical and bioanalytical analyses.

Immunoblot analysis of tumor lysates revealed marked suppression of AKT phosphorylation following treatment with all three compounds (Figure 3A–C). GT220 and AZD8186 produced dose-dependent inhibition of AKT phosphorylation at both Ser473 and Thr308, demonstrating effective pathway suppression across the tested dose range (Figure 3A–C). In contrast, KIN193 showed less pronounced inhibition (Figure 3A–C).

**Figure 3.**
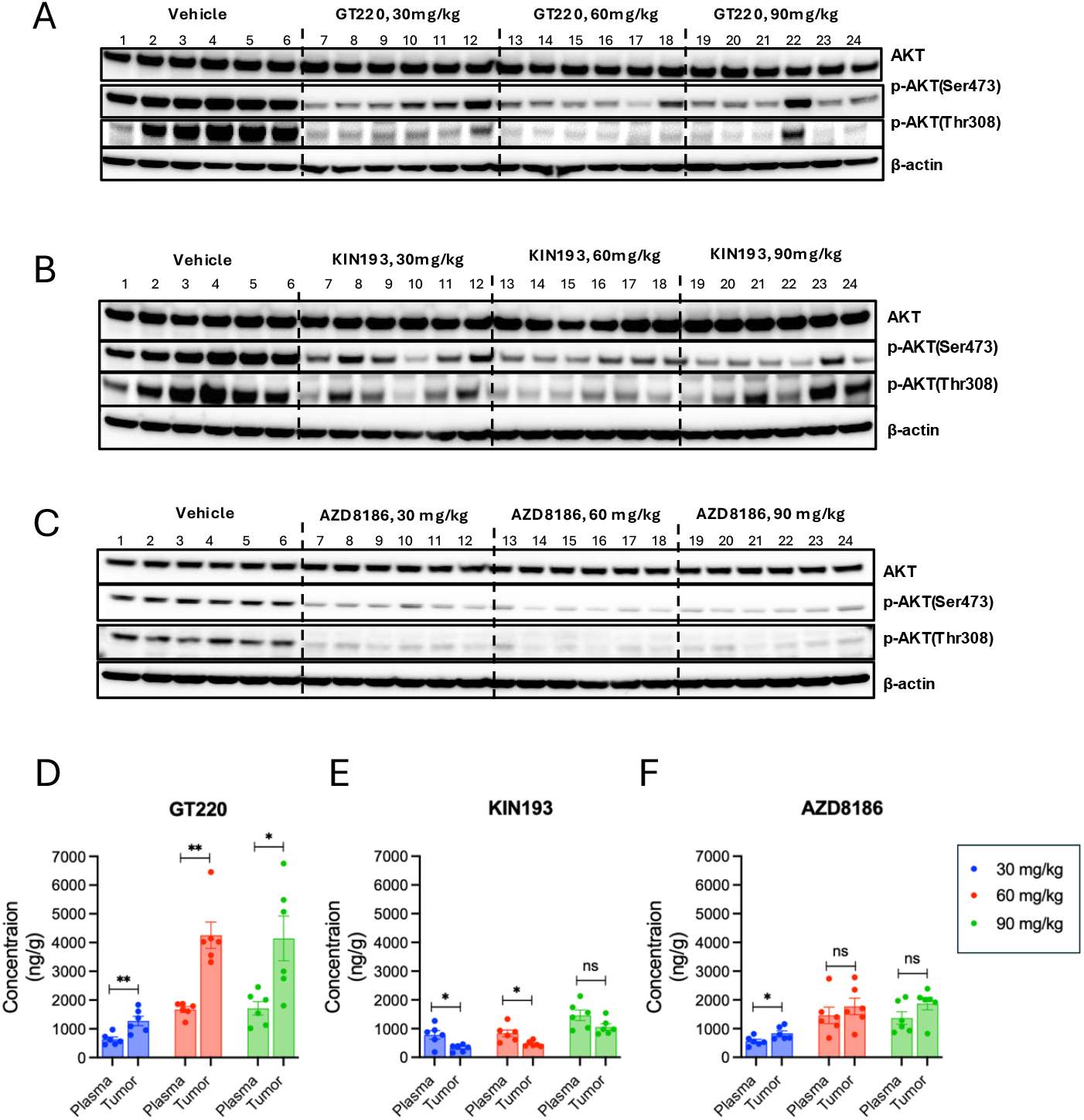
Pharmacodynamic (PD) and pharmacokinetic (PK) characterization of GT220 and other PI3Kβ inhibitors in PTEN-deficient HCC70 xenografts. BALB/c nude mice bearing HCC70 xenografts were treated with GT220, KIN193, or AZD8186 at doses of 30, 60, or 90 mg/kg, administered orally twice daily (po, bid; n = 6 per group) for 3 days. Tumors and plasma were collected 1.5 hours after the final dose. **(A–C)** Tumor lysates were analyzed by immunoblotting for phosphorylated AKT (p-AKT) levels using the indicated antibodies. **(D-F)** Intratumoral and corresponding plasma drug concentrations measured by LC–MS/MS at the indicated doses (mg/kg) for each compound. Data in (**D**-**F**) are presented as mean ± SEM, with individual data points shown. Statistical significance is indicated (*P < 0.05; **P < 0.01).

Pharmacokinetic analysis by LC–MS/MS demonstrated that GT220 achieved substantially higher intratumoral drug concentrations relative to matched plasma compared with KIN193 and AZD8186 (Figure 3D–F). Notably, GT220 achieved the highest tumor exposure among the compounds tested and exhibited approximately two-fold higher concentrations in tumor tissue than in plasma across doses, whereas KIN193 and AZD8186 showed tumor concentrations that were comparable to or lower than plasma levels (Figure 3D-F). These pharmacokinetic properties correlated with the robust pharmacodynamic suppression of AKT signaling observed in GT220-treated tumors.

Collectively, these results indicate that GT220 achieves favorable tumor exposure and sustained PI3Kβ pathway inhibition in vivo, consistent with its superior antitumor activity in PTEN-deficient HCC70 xenograft models.

### GT220 demonstrates robust antitumor efficacy and favorable tolerability in PTEN-deficient HCC70 xenografts

To evaluate the in vivo antitumor efficacy of GT220, BALB/c nude mice bearing established PTEN-deficient HCC70 xenografts were treated orally twice daily with GT220 at doses of 30, 60, or 90 mg/kg for 21 days. GT220 produced a dose-dependent suppression of tumor growth, with significant tumor inhibition observed across all doses tested (Figure 4A). Notably, substantial antitumor activity was achieved at the lowest dose of 30 mg/kg, indicating strong in vivo potency.

**Figure 4.**
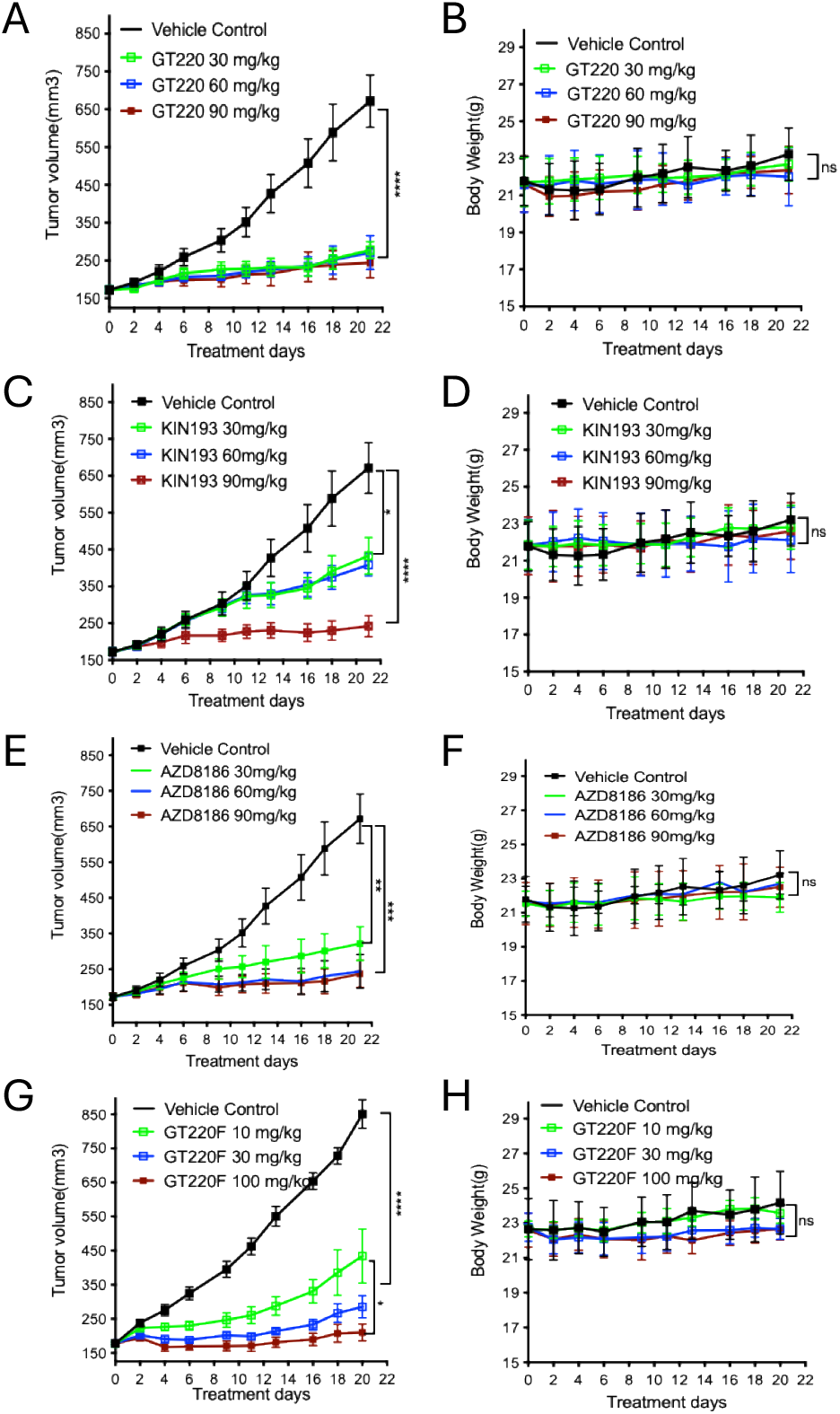
In vivo antitumor efficacy and tolerability of GT220 in the HCC70 xenograft model. BALB/c nude mice bearing established HCC70 (PTEN-deficient) xenografts were treated orally twice daily (po, bid; n = 6 per group) with GT220 (**A, B**), KIN193 (**C. D**), or AZD8186 (**E, F**) at doses of 30, 60, or 90 mg/kg for 21 days. Tumor volumes (**A, C, E**) and body weights (B, D, F) were measured every 2–3 days throughout the treatment period. (**G, H**) Antitumor efficacy and tolerability of the salt form GT220F administered orally twice daily at 10, 30, or 100 mg/kg (n = 6 per group) for 21 days. Tumor volumes (**G**) and body weights **(H**) were monitored longitudinally. Data are presented as mean ± SEM. Statistical significance is indicated as shown (P < 0.05, *P < 0.01, **P < 0.005, ***P < 0.0001; one-way ANOVA). “ns” denotes not significant.

For comparison, mice bearing HCC70 xenografts were treated in parallel with the PI3Kβ-selective inhibitor KIN193 or the dual PI3Kβ/δ inhibitor AZD8186 using identical dosing regimens. Although both comparators exhibited antitumor activity, GT220 consistently demonstrated equal or greater tumor growth inhibition across dose levels (Figure 4A, C, E).

Throughout the 21-day treatment period, body weights remained stable in all treatment groups, including those receiving the highest doses of GT220, KIN193, or AZD8186 (Figure 4B, D, F), indicating good tolerability and an absence of overt toxicity.

To further assess formulation-dependent efficacy, the salt form of GT220 (GT220F) was evaluated in a separate cohort of HCC70 xenograft–bearing mice. Oral administration of GT220F at doses of 10, 30, or 100 mg/kg twice daily for 21 days resulted in dose-dependent tumor growth inhibition, with significant efficacy observed at 30 and 100 mg/kg (Figure 4G). Body weights remained stable across all GT220F dose groups (Figure 4H), further supporting the favorable tolerability of GT220 formulations.

Collectively, these data demonstrate that GT220 exhibits robust and sustained antitumor efficacy in PTEN-deficient HCC70 xenografts, coupled with good tolerability across a broad dose range, supporting its continued development as a PI3Kβ-targeted therapeutic.

### GT220 lacks antitumor efficacy and pathway inhibition in PTEN–wild-type HCC1954 xenografts

To determine whether the in vivo activity of GT220 is restricted to PI3Kβ-dependent tumors, we evaluated its antitumor efficacy in the PTEN–wild-type HCC1954 xenograft model, which harbors an activating *PIK3CA* mutation and relies predominantly on PI3Kα signaling. BALB/c nude mice bearing established HCC1954 xenografts were treated orally twice daily with vehicle or GT220 at doses of 60 or 90 mg/kg for 16 days.

GT220 treatment did not significantly affect tumor growth at either dose compared with vehicle control (Figure 5A). Consistent with the lack of antitumor activity, body weights remained stable across all treatment groups throughout the dosing period, indicating good tolerability but no evident therapeutic effect in this model (Figure 5B).

**Figure 5.**
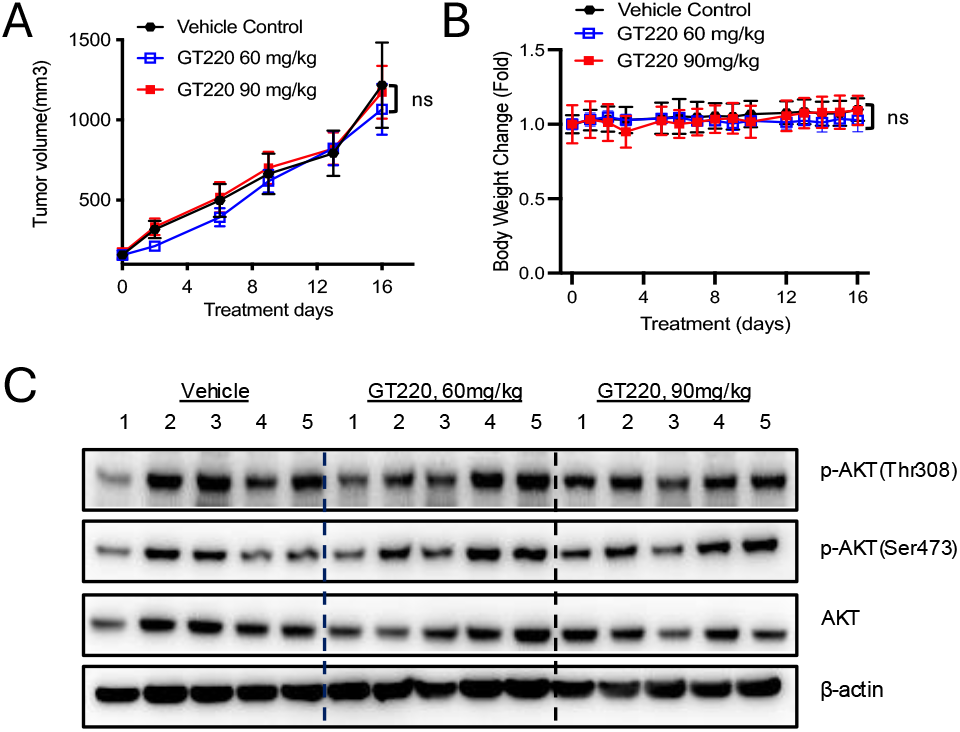
Lack of antitumor efficacy and pathway inhibition by GT220 in PTEN-wild-type HCC1954 xenografts. **(A–B)** BALB/c nude mice bearing HCC1954 (PTEN-wild-type) xenografts were treated with vehicle or GT220 at 60 or 90 mg/kg, administered orally twice daily (po, bid; n = 5 per group) for 16 days. Tumor volumes and body-weight changes (**B**) were monitored throughout the treatment period. No statistically significant differences were observed relative to vehicle control (ns, one-way ANOVA). (**C**) In a separate PD cohort, HCC1954 xenograft–bearing mice (n = 5 per group) were treated with vehicle or GT220 (60 or 90 mg/kg, po, bid) for 3.5 days. Tumors were harvested 1.5 hours after the final dose, and AKT phosphorylation at Thr308 and Ser473 was assessed by immunoblotting, with total AKT and β-actin serving as controls.

To assess whether the absence of efficacy reflected insufficient pathway inhibition, a separate pharmacodynamic cohort was treated with vehicle or GT220 (60 or 90 mg/kg, po, bid) for 3.5 days. Tumors were harvested 1.5 hours after the final dose and analyzed for AKT phosphorylation. Immunoblotting revealed that GT220 failed to suppress AKT phosphorylation at either Thr308 or Ser473 in HCC1954 tumors (Figure 5C), despite adequate dosing.

Together, these results demonstrate that GT220 does not inhibit PI3K signaling or tumor growth in PTEN–wild-type, PI3Kα-dependent HCC1954 xenografts, underscoring the context-dependent activity of GT220 and reinforcing its selectivity for PI3Kβ-driven tumors.

## Discussion

In this study, we describe the discovery and preclinical characterization of GT220, a next-generation, highly selective PI3Kβ inhibitor designed to target cancers driven by *PIK3CB* alterations or PTEN loss. Our data demonstrate that GT220 exhibits exceptional biochemical and cellular selectivity for PI3Kβ, potently suppresses AKT signaling and cell viability in PTEN-deficient and *PIK3CB*-mutant cancer cells, and achieves robust antitumor activity in vivo in PTEN-deficient xenograft models. Importantly, GT220 lacks activity in PTEN–wild-type, PI3Kα-dependent models both in vitro and in vivo. Together, these findings provide compelling evidence that selective targeting of PI3Kβ represents a viable therapeutic strategy for a defined subset of cancers.

Although the PI3K pathway has been extensively pursued as a therapeutic target, most clinical efforts have focused on PI3Kα and PI3Kδ. These strategies have yielded important successes, including the approval of alpelisib for *PIK3CA*-mutant breast cancer and idelalisib for B-cell malignancies. However, the therapeutic windows of these agents are constrained by the essential roles of PI3Kα and PI3Kδ in normal physiology, resulting in dose-limiting toxicities and limited tolerability. In contrast, PI3Kβ occupies a unique biological niche. Accumulating genetic and functional evidence—including foundational work from our laboratory—has established PI3Kβ as a critical driver of oncogenic signaling in tumors with PTEN loss or activating *PIK3CB* mutations, while appearing largely dispensable in most adult tissues. This differential dependency suggests that selective PI3Kβ inhibition may achieve more favorable efficacy– toxicity profiles than inhibitors targeting other PI3K isoforms.

Despite this strong rationale, prior attempts to therapeutically target PI3Kβ have faced substantial challenges. Clinical-stage PI3Kβ inhibitors such as GSK2636771 and AZD8186 have been limited by suboptimal selectivity and pharmacologic properties. AZD8186, a dual PI3Kβ/δ inhibitor, was discontinued due to dose-limiting toxicities likely driven by unintended PI3Kδ inhibition. Similarly, although GSK2636771 has been described as PI3Kβ-selective, our biochemical analyses reveal comparable inhibitory activity against PI3Kβ and PI3Kδ. Notably, plasma concentrations achieved in early-phase clinical studies exceeded the IC_50_ values for both isoforms, providing a plausible explanation for the gastrointestinal and metabolic toxicities observed in patients. These experiences underscore a central challenge in PI3Kβ drug development: achieving sufficient potency against PI3Kβ while avoiding off-target inhibition of PI3Kδ and other kinases.

GT220 addresses several of these limitations. Through an integrated AI-driven design and medicinal chemistry optimization scheme, GT220 was engineered to achieve high affinity and exceptional selectivity for PI3Kβ across the kinome. In contrast to prior compounds, GT220 exhibits minimal activity against PI3Kδ, translating into selective suppression of PI3Kβ-driven signaling and tumor growth. In vivo, GT220 demonstrates favorable pharmacokinetic properties, including enhanced tumor exposure relative to plasma and sustained inhibition of AKT phosphorylation in PTEN-deficient tumors. These features correlate with robust antitumor efficacy in both breast and prostate PTEN-null xenograft models, while sparing PI3Kα-dependent tumors. Collectively, these attributes suggest that GT220 may offer a substantially improved therapeutic index compared with earlier PI3Kβ inhibitors.

Beyond PTEN loss, our findings also have implications for cancers harboring activating *PIK3CB* mutations. Although less frequent than *PIK3CA* alterations, *PIK3CB* mutations are recurrent across multiple tumor types and remain largely unaddressed therapeutically. The selective activity of GT220 in *PIK3CB*-mutant models supports the concept that these alterations define a distinct, actionable molecular subset of cancers that may benefit from PI3Kβ-directed therapy. Future studies will be important to further define predictive biomarkers, resistance mechanisms, and optimal combination strategies—particularly in the context of immune modulation, given emerging roles of PI3Kβ in tumor–immune interactions.

In summary, our work establishes GT220 as a highly selective and potent PI3Kβ inhibitor with strong preclinical efficacy in PI3Kβ-dependent cancers driven by PTEN loss or *PIK3CB* mutations. These findings not only validate PI3Kβ as an actionable oncogenic target, but also highlight the importance of isoform selectivity in achieving meaningful therapeutic benefit without off-target risks. GT220 represents a promising candidate for clinical development and provides a strong foundation for precision targeting of PI3Kβ-driven malignancies.

## Materials and methods

### Compounds and Reagents

GT-220 was synthesized by WuXi AppTec. KIN193, AZD8186, GSK2636771, BYL719 were purchased from MedChemExpress. For *in vitro* studies, compounds were dissolved in DMSO. For *in vivo* studies, compounds were dissolved in 10% DMSO+40% PEG300+5%Tween80+45% Saline and administered by oral gavage at 30, 60 or 90 mg/kg twice a day.

### KINOMEscan analysis

Kinase profiling was conducted by DiscoverX (Eurofins DiscoverX Corporation, San Diego, CA, USA) using the KINOMEscan scanMAX lead hunter panel. The panel employs a biochemical assay to measure drug binding utilizing a panel of 468 DNA-tagged kinases. The tested compound was evaluated at a concentration of 10 mM. Compounds binding to the kinase active site reduce the amount of kinase bound to a solid support with an immobilized ligand. Quantification is done using qPCR to detect the associated DNA label. The strength of compound binding is assessed based on its interference with kinase-ligand binding, and results are reported as percent control (%CTRL), indicating the percentage of kinase remaining bound to the immobilized ligand.

### Dissociation Constant assay

Binding constants (Kds) of PIK3CA, PIK3CB, PIK3CD, and PIK3CG were evaluated by DiscoverX (Eurofins DiscoverX Corporation, San Diego, CA, USA) using the KINOMEscan KdELECT lead hunter panel. The assay involves three components: a DNA-tagged kinase, an immobilized ligand, and a test compound. The competition ability of the test compound is measured via quantitative PCR of the DNA tag. Test compounds were prepared as 111X stocks in 100% DMSO. Binding constants (Kds) were determined using an 11-point, 3-fold dilution series with three DMSO control points. Compounds for Kd measurements were distributed by acoustic transfer (non-contact dispensing) in 100% DMSO. Kds were calculated with a standard dose-response curve using the Hill equation: Response = Background +(Signal – Background/ (1 + (Kd ^Hill Slope^ / Dose ^Hill Slope^)) The Hill Slope was set to -1.

### Cell Culture

HCC70 and HCC1954 were purchase from ATCC and maintained *in vitro* as a monolayer culture in RPMI1640, DMEM or EMEM supplemented with 10% heat inactivated fetal bovine serum and 1% Pen-Strep at 37ºC in an atmosphere of 5% CO_2_ in air. The cells were routinely sub-cultured twice weekly by trypsin-EDTA treatment. The cells growing in an exponential growth phase were harvested and counted for tumor inoculation.

### Cell viability assay

Cells were seeded in 96-well plates at a density of 2,500 cells per well and treated with ten-fold serial dilutions of compounds with a starting concentration of 10 μM. Cell viability was assessed after three days of treatment by CellTiter-Glo (Promega). Curve fitting analysis and IC_50_ value determination were performed using GraphPad Prism 10.

### Western blot analysis

Western blot analysis was performed as described previously [30, 31]. Anti-phospho-AKT(Thr308) (#2965S), Anti-phospho-AKT (Ser 473) (#4060S), Anti-AKT(#9272S), antibodies were purchased from Cell Signaling Technology. Anti-Vinculin antibody was purchased from Sigma.

### Antitumor experiments

Animal experiments of HCC70 and HCC2954 breast cancer xenografts were performed at WuXi AppTec according to the local regulations.

The HCC70 tumor cells will be maintained *in vitro* as a monolayer culture in RPMI1640 supplemented with 10% heat inactivated fetal bovine serum and 1% Antibiotic-Antimycotic, at 37ºC in an atmosphere of 5% CO_2_ in air. BALB/c nude, female, 6∼8 weeks, weighing approximately 18-22g. Each mouse will be inoculated subcutaneously at the right upper flank with HCC70 tumor cells (10 × 10^6^) in 0.2 mL of PBS with matrigel (1:1) for tumor development.

The tumor-bearing animals were randomized, and treatment was started when the average tumor volume reaches approximately 150 mm^3^ for the efficacy study. In the cohort of mice with HCC70 xenograft tumors, GT-220, KIN193, AZD8186 were administrated at 30, 60, 90 mg/kg twice a day. In the cohort of mice with PC3 xenograft tumors, GT-220 (b.i.d) and GSK2636771(q.d.) were administrated at 3, 10, 30 mg/kg. Tumor sizes and body weight will be measured three times per week in two dimensions using a caliper, and the volume will be expressed in mm^3^ using the formula: V = 0.5 *a* x *b*^2^ where *a* and *b* are the long and short diameters of the tumor, respectively.

### Pharmacodynamic studies

When the average tumor size reached approximately 0.4 cm3 in HCC70, tumors will be harvested at 1.5 hours after the last dose and divided into two sections. Both sections will be rapidly frozen in liquid nitrogen. One section will be utilized for exposure determination at WuXi, while the other section will be used for Western blotting analysis, including the examination of phospho-AKT (Ser473) (CST #4060), phospho-AKT (Thr308) (CST #2965), total AKT (CST #9272), and β-actin. Plasma samples collected at 1.5 hours post the last dose will be stored at - 80ºC for pharmacokinetic (PK) analysis at WuXi AppTec.

## Statistical analysis

Statistical significance was determined using unpaired Student’s *t*-tests or ANOVA by GraphPadPrism 10 (GraphPad Software). Data are considered significant when *P* values are < 0.05.

## Reference

1. Liu, P., et al., Targeting the phosphoinositide 3-kinase pathway in cancer. Nat Rev Drug Discov, 2009. 8(8): p. 627–44.

2. Thorpe, L.M., H. Yuzugullu, and J.J. Zhao, PI3K in cancer: divergent roles of isoforms, modes of activation and therapeutic targeting. Nat Rev Cancer, 2015. 15(1): p. 7–24.

3. Samuels, Y., et al., High frequency of mutations of the PIK3CA gene in human cancers. Science, 2004. 304(5670): p. 554.

4. Lawrence, M.S., et al., Discovery and saturation analysis of cancer genes across 21 tumour types. Nature, 2014. 505(7484): p. 495–501.

5. Andre, F., et al., Alpelisib for PIK3CA-Mutated, Hormone Receptor-Positive Advanced Breast Cancer. N Engl J Med, 2019. 380(20): p. 1929–1940.

6. Andre, F., et al., Alpelisib plus fulvestrant for PIK3CA-mutated, hormone receptor-positive, human epidermal growth factor receptor-2-negative advanced breast cancer: final overall survival results from SOLAR-1. Ann Oncol, 2021. 32(2): p. 208–217.

7. Whale, A.D., et al., Functional characterization of a novel somatic oncogenic mutation of PIK3CB. Signal Transduct Target Ther, 2017. 2: p. 17063.

8. Karlsson, T., et al., Endometrial cancer cells exhibit high expression of p110beta and its selective inhibition induces variable responses on PI3K signaling, cell survival and proliferation. Oncotarget, 2017. 8(3): p. 3881–3894.

9. Nakanishi, Y., et al., Activating Mutations in PIK3CB Confer Resistance to PI3K Inhibition and Define a Novel Oncogenic Role for p110beta. Cancer Res, 2016. 76(5): p. 1193–203.

10. Fusco, N., et al., PTEN Alterations and Their Role in Cancer Management: Are We Making Headway on Precision Medicine? Genes (Basel), 2020. 11(7).

11. Milella, M., et al., PTEN: Multiple Functions in Human Malignant Tumors. Front Oncol, 2015. 5: p. 24.

12. Ma, C.X., et al., A Phase I Trial of BKM120 (Buparlisib) in Combination with Fulvestrant in Postmenopausal Women with Estrogen Receptor-Positive Metastatic Breast Cancer. Clin Cancer Res, 2016. 22(7): p. 1583–91.

13. Garrido-Castro, A.C., et al., Phase 2 study of buparlisib (BKM120), a pan-class I PI3K inhibitor, in patients with metastatic triple-negative breast cancer. Breast Cancer Res, 2020. 22(1): p. 120.

14. Juric, D., et al., Phosphatidylinositol 3-Kinase alpha-Selective Inhibition With Alpelisib (BYL719) in PIK3CA-Altered Solid Tumors: Results From the First-in-Human Study. J Clin Oncol, 2018. 36(13): p. 1291–1299.

15. Jia, S., et al., Essential roles of PI(3)K-p110beta in cell growth, metabolism and tumorigenesis. Nature, 2008. 454(7205): p. 776–9.

16. Liu, P., et al., Oncogenic PIK3CA-driven mammary tumors frequently recur via PI3K pathway-dependent and PI3K pathway-independent mechanisms. Nat Med, 2011. 17(9): p. 1116–20.

17. Utermark, T., et al., The p110alpha and p110beta isoforms of PI3K play divergent roles in mammary gland development and tumorigenesis. Genes Dev, 2012. 26(14): p. 1573–86.

18. Wang, Q., et al., PI3K-p110alpha mediates resistance to HER2-targeted therapy in HER2+, PTEN-deficient breast cancers. Oncogene, 2016. 35(27): p. 3607–12.

19. Yuzugullu, H., et al., A PI3K p110beta-Rac signalling loop mediates Pten-loss-induced perturbation of haematopoiesis and leukaemogenesis. Nat Commun, 2015. 6: p. 8501.

20. Wee, S., et al., PTEN-deficient cancers depend on PIK3CB. Proc Natl Acad Sci U S A, 2008. 105(35): p. 13057–62.

21. Schwartz, S., et al., Feedback suppression of PI3Kalpha signaling in PTEN-mutated tumors is relieved by selective inhibition of PI3Kbeta. Cancer Cell, 2015. 27(1): p. 109–22.

22. Peng, W., et al., Loss of PTEN Promotes Resistance to T Cell-Mediated Immunotherapy. Cancer Discov, 2016. 6(2): p. 202–16.

23. Juric, D., et al., Convergent loss of PTEN leads to clinical resistance to a PI(3)Kalpha inhibitor. Nature, 2015. 518(7538): p. 240–4.

24. Bergholz, J.S., et al., PI3Kbeta controls immune evasion in PTEN-deficient breast tumours. Nature, 2023. 617(7959): p. 139–146.

25. Ni, J., et al., Functional characterization of an isoform-selective inhibitor of PI3K-p110beta as a potential anticancer agent. Cancer Discov, 2012. 2(5): p. 425–33.

26. Bedard, P.L., et al., First-in-human trial of the PI3Kbeta-selective inhibitor SAR260301 in patients with advanced solid tumors. Cancer, 2018. 124(2): p. 315–324.

27. Mateo, J., et al., A First-Time-in-Human Study of GSK2636771, a Phosphoinositide 3 Kinase Beta-Selective Inhibitor, in Patients with Advanced Solid Tumors. Clin Cancer Res, 2017. 23(19): p. 5981–5992.

28. Choudhury, A.D., et al., A Phase I Study Investigating AZD8186, a Potent and Selective Inhibitor of PI3Kbeta/delta, in Patients with Advanced Solid Tumors. Clin Cancer Res, 2022. 28(11): p. 2257–2269.

29. Barlaam, B., et al., Discovery of (R)-8-(1-(3,5-difluorophenylamino)ethyl)-N,N-dimethyl-2-morpholino-4-oxo-4H-chromene-6-carboxamide (AZD8186): a potent and selective inhibitor of PI3Kbeta and PI3Kdelta for the treatment of PTEN-deficient cancers. J Med Chem, 2015. 58(2): p. 943–62.

30. Sun, B., et al., Inhibition of the transcriptional kinase CDK7 overcomes therapeutic resistance in HER2-positive breast cancers. Oncogene, 2020. 39(1): p. 50–63.

31. Wang, Q., E. Weisberg, and J.J. Zhao, The gene dosage of class Ia PI3K dictates the development of PTEN hamartoma tumor syndrome. Cell Cycle, 2013. 12(23): p. 3589–93.

